# Not a single clock: multiple behavioral rhythms in perceptual averaging

**DOI:** 10.64898/2026.08.19.745670

**Authors:** Maëlan Q. Menétrey, David Pascucci

## Abstract

Several theories propose that perception and attention are governed by rhythmic processes that give rise to periodic fluctuations in behavior. However, empirical support for behavioral rhythms has been derived largely from paradigms involving brief, static stimuli. Here, we introduce a temporal averaging task requiring integration of rapidly unfolding visual features. Across three experiments, we tested averaging of orientation, size, and color under different eccentricity conditions. We used a temporally weighted averaging model to assess whether the influence of individual stimulus samples on perceptual estimates exhibits periodic modulation over time. We found no common rhythmic signature across tasks. Instead, orientation and size judgments showed reliable low-frequency modulations (<2.5 Hz), whereas color judgments showed only weak trends. Higher-frequency components (∼3.5–8 Hz), often linked to theta and alpha rhythms, were observed only in a subset of participants and were limited to parafoveal orientation processing. These findings challenge the notion of universal behavioral rhythms and instead suggest that temporal dynamics are task-dependent, with slow oscillatory processes emerging as the most consistent feature.

## Introduction

Recent research has proposed that perceptual sensitivity and spatial attention are modulated by rhythmic fluctuations. This has led to theories such as *perceptual cycles* (Samaha & Postle, 2015; VanRullen, 2016; Kaltenmaier et al., 2025) and *rhythmic theories of attention* (Landau & Fries, 2012; Helfrich et al., 2018; Fiebelkorn & Kastner, 2019), which suggest a hierarchy of intrinsic rhythms in human performance, with slower rhythms in the theta band (4–7 Hz) modulating attentional sampling in space and faster ones in the alpha band (8–13 Hz) affecting the detection and discrimination of local stimuli (Fries, 2015; Bonnefond et al., 2024). This has been accompanied by direct evidence of these *behavioral rhythms* in perceptual and attentional tasks (Fiebelkorn et al., 2013; Song et al., 2014; Drewes et al., 2015; Huang et al., 2015).

In behavioral studies, rhythms are typically assessed using dense-sampling paradigms, in which the onset of a target stimulus is systematically varied relative to a reference, or ‘resetting’, event. This resetting event, typically a discrete sensory signal (e.g., a visual flash or auditory tone), serves as a temporal reference that is supposed to align ongoing fluctuations in performance across trials, allowing behavioral responses to be reconstructed as a function of time relative to that event (Landau & Fries, 2012). For example, in spatial attention, rhythmic fluctuations in detection performance can be assessed using cue–target paradigms, where targets appear at varying delays after a spatial cue (e.g., Chota et al., 2022). Similarly, in perceptual detection or discrimination tasks, performance is sampled across variable stimulus onset asynchronies relative to a reference event (e.g., Michel et al., 2021). In all cases, the presence of rhythms is inferred from the spectral content of performance fluctuations, time-locked to the resetting event and aggregated across many trials (but see Brookshire, 2022).

Evidence for the existence and robustness of behavioral rhythms, however, remains mixed and is largely confined to detection and discrimination paradigms using briefly flashed stimuli, isolated from natural temporal and spatial structure (for a review, see Kienitz et al., 2021). This leaves several important questions open. First, it is unclear whether behavioral rhythms generalize to more dynamic stimulus conditions in which visual events unfold continuously over time and space, rather than appearing as discrete, localized transients (Pascucci & Kristjánsson, 2026). Second, it is unclear whether they reflect general or distinct oscillatory dynamics in specific visual circuits, potentially giving rise to feature-dependent temporal structure across partially segregated processing streams (e.g., orientation, color, and size). Third, it is unclear whether these effects depend on the spatial structure of visual input, for instance whether they vary with eccentricity.

To address these questions, we tested for periodic components in how human observers extract average features from rapid spatiotemporal sequences of stimuli. We used an extension of a weighted-average model (Anderson, 1967; Pascucci et al., 2021; Tiurina et al., 2024), adapted to the temporal domain. In three experiments, participants reproduced the perceived average orientation (Experiment 1, Figure 1A), size (Experiment 2, Figure 2A), or color (Experiment 3, Figure 3A) of a rapid stream of stimuli, and their responses were modeled to estimate the temporal weights with which each stimulus sample influenced the final, reported percept. Stimuli were presented at two retinal eccentricities to assess whether rhythmic components vary as a function of spatial location and on the spacing between successive stimulus samples. To quantify behavioral rhythms, we applied spectral analysis to the temporal weighting profiles, separately for each experiment (orientation, size, or color) and eccentricity condition. Our results revealed feature- and eccentricity-dependent rhythmic components that were not consistently observed across conditions and were largely driven by in-phase fluctuations in temporal weights across participants, with an overall dominance of slow rhythms (<2.5 Hz).

Overall, these results point to a heterogeneous pattern, indicating that behavioral rhythmicity is not a stable, general property of human performance but varies across features and experimental contexts.

## Results

Three independent groups of participants reproduced the perceived average of a rapidly presented sequence of visual features: orientation (Experiment 1, N = 22), size (Experiment 2, N = 21), and color (Experiment 3, N = 25). Stimuli were presented at two eccentricities (parafovea: 3.5°, periphery: 7°) and updated every 25 ms (40 Hz).

To characterize the contribution of individual stimulus samples (i.e., successive feature presentations within the sequence) to perceptual judgments, we modeled reproduction errors as a function of the feature values presented at each temporal position using a weighted-averaging model adapted to the temporal domain (Anderson, 1967; Pascucci et al., 2021; Tiurina et al., 2024). In this model, the reproduction error on each trial is expressed as a linear combination of the deviations of individual samples from the sequence mean. Model parameters were estimated using least squares, yielding a temporal weighting profile that quantifies the relative influence of each sample on the final perceptual estimate. This approach makes it possible to identify systematic biases in perceptual averaging, for example, whether samples presented at specific moments are over- or under-weighted, and to test for periodic fluctuations in these weights. Such fluctuations would indicate oscillatory structure in the temporal dynamics of perceptual averaging. We therefore subjected the temporal weighting profiles to spectral analysis, separately for each experiment and eccentricity condition (see Methods).

In Experiment 1, participants reproduced the perceived average orientation of a sequence of Gabor patches (Figure 1A) by adjusting the orientation of a response tool consisting of two dots connected by an imaginary line (Ceylan et al., 2021; Ozkirli & Pascucci, 2023). After cleaning outlier responses (Methods and Supplementary Figure S1A), we quantified performance using the circular standard deviation of the response errors (parafoveal: 10.88 ± 0.57; peripheral: 10.78 ± 0.6; no difference between conditions: *t*(21) = 0.51, *p* = .61; Supplementary Figure S1B). Spectral analysis of the temporal weighting profiles revealed systematic oscillatory components in the parafoveal condition, with significant peaks at 1.5, 3.75, and 7.5 Hz (all *p*s < .05 with FDR corrected, only the two higher frequencies survived cluster-based permutation; Figures 1B and 1C). At these frequencies, we observed reliable phase clustering across participants (Rayleigh tests: all *p*s < .003; Figure 1B), indicating consistent temporal alignment in the weighting profile. In contrast, the periphery condition exhibited a simpler pattern, with a single low-frequency component (peak = 1.25 Hz, *p* < .05 both FDR-corrected and cluster-based permutation tests; Rayleigh test: *p* = < .001; Figures 1B and 1C). These patterns were not uniformly expressed across individuals: only 27.3% of participants exhibited strong phase alignment with the group mean phase across all significant peaks observed at the group level, with substantial variability across frequencies and eccentricities (Supplementary Figure S2A), and over time (Supplementary Figure S2B).

**Figure 1.**
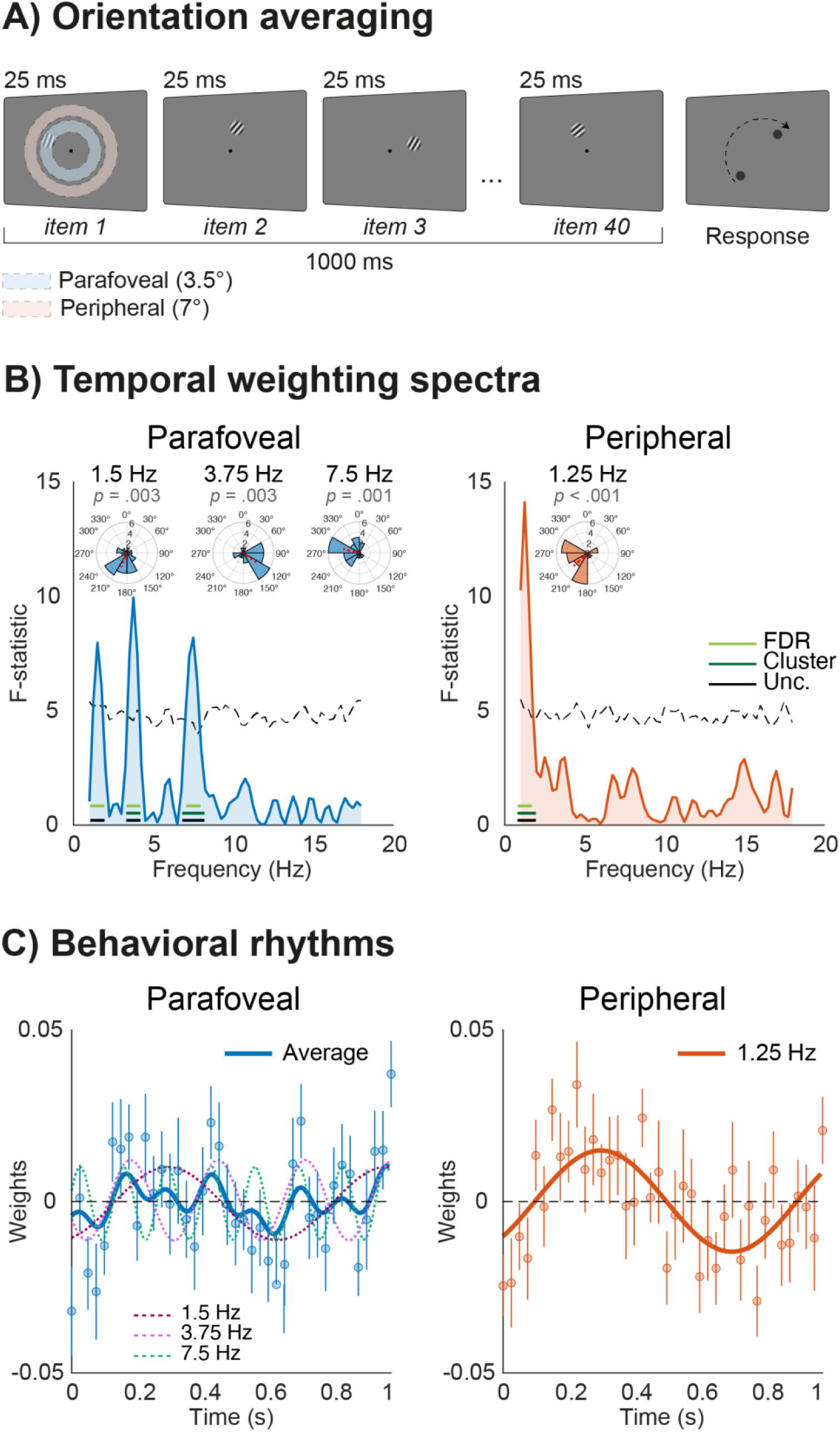
Orientation judgments. A) In Experiment 1, participants viewed a stream of 40 Gabors presented at 25-ms intervals (40 Hz), either at parafoveal or peripheral eccentricities, and reported the perceived average orientation by means of a reproduction response. B) Spectral analysis of the temporal weighting profiles in parafoveal (left panel) and peripheral (right panel) conditions. Significant peaks (*p* < .05) are shown for uncorrected (black lines; GLM F-statistic), FDR-corrected (light green lines, GLM F-statistic corrected using the Benjamini–Hochberg procedure), and cluster-based permutation tests (dark green lines; cluster mass computed as the sum of the F-statistic within each cluster, see Methods). For visualization, the dashed line indicates the 99th percentile of the surrogate distribution. The polar plots show phase clustering for the FDR-corrected significant peaks, along with the corresponding Rayleigh test p-values. Histograms show the distribution of individual participant phases; the red line indicates the circular mean phase (resultant vector across participants). C) Grand-averaged time course of the temporal weighting profile in parafoveal (left panel) and peripheral (right panel) conditions. Blue and orange lines represent the mean of FDR-corrected significant rhythms in the parafoveal condition, where multiple rhythms are present, and the single identified rhythm in the peripheral condition, respectively. Individual FDR-corrected significant rhythms are also shown for the parafoveal condition. Dots and error bars indicate the mean at each time point and the corresponding standard error of the mean (SEM).

In Experiment 2, participants reproduced the perceived average size of a sequence of light gray filled spots (Figure 2A). After cleaning outlier responses (Methods and Supplementary Figure S1A), we quantified performance using the standard deviation of the response errors (parafoveal: 13.21 ± 0.65; peripheral: 14.57 ± 0.75; Supplementary Figure S1B). For this experiment, a significant difference between eccentricities was found (*t*(20) = 2.5, *p* = .02), suggesting that the task was more difficult in the peripheral condition. Despite this difference, spectral analysis of the temporal weighting profiles revealed a single low-frequency component in both parafoveal and peripheral conditions, contrasting with the more complex spectral profile observed for orientation judgments in the parafoveal condition of Experiment 1. Significant peaks were found at 1.25 Hz and 1 Hz in the parafoveal and peripheral conditions, respectively (all *p*s < .05 with FDR corrected, only the peak in parafoveal condition survived cluster-based permutation; Figures 2B and 2C), with reliable phase clustering across participants within each condition (Rayleigh tests: all *p*s < .01; Figure 2B). As in Experiment 1, however, these effects were not consistently expressed across individuals, with only 33% of participants showing strong alignment with the group mean phase at the relevant peak frequencies in both conditions (Supplementary Figure S2A). There was also substantial variability over time (Supplementary Figure S2B).

**Figure 2.**
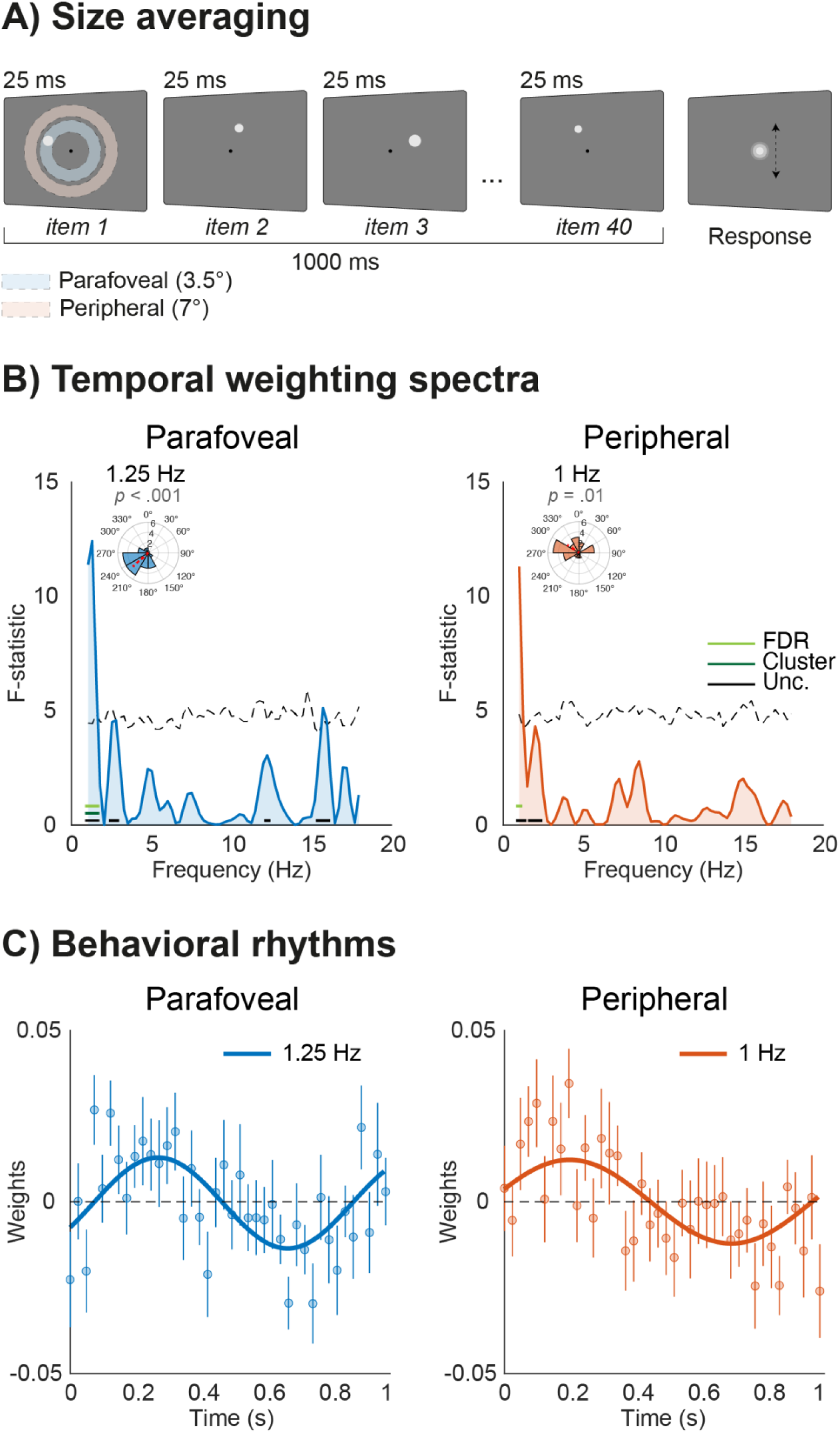
Size judgments. A) In Experiment 2, participants viewed a stream of 40 gray spots presented at 25-ms intervals (40 Hz), either at parafoveal or peripheral eccentricities, and reported the perceived average size by means of a reproduction response. B) Spectral analysis of the temporal weighting profiles in parafoveal (left panel) and peripheral (right panel) conditions. Significant peaks (*p* < .05) are shown for uncorrected (black lines; GLM F-statistic), FDR-corrected (light green lines, GLM F-statistic corrected using the Benjamini– Hochberg procedure), and cluster-based permutation tests (dark green lines; cluster mass computed as the sum of the F-statistic within each cluster, see Methods). For visualization, the dashed line indicates the 99th percentile of the surrogate distribution. The polar plots show phase clustering for the FDR-corrected significant peaks, along with the corresponding Rayleigh test p-values. Histograms show the distribution of individual participant phases; the red line indicates the circular mean phase (resultant vector across participants). C) Grand-averaged time course of the temporal weighting profile in parafoveal (left panel) and peripheral (right panel) conditions. Blue and orange lines represent the single FDR-corrected significant rhythms in parafoveal and peripheral conditions, respectively. Dots and error bars indicate the mean at each time point and the corresponding standard error of the mean (SEM).

In Experiment 3, participants reproduced the perceived average color of a sequence of colored spots (Figure 3A). As in Experiments 1, we cleaned outlier responses (Methods and Supplementary Figure S1A) and quantified performance using the circular standard deviation of the response errors (parafoveal: 20.23 ± 0.75; peripheral: 20.14 ± 0.68; no difference between conditions: *t*(24) = 0.27, *p* = .78; Supplementary Figure S1B). Unlike Experiments 1 and 2, however, spectral analysis of the temporal weighting profiles did not reveal any oscillatory components that survived correction for multiple comparisons (FDR correction and cluster-based permutation) in either the parafoveal or peripheral condition (Figure 3B). Only week rhythmic modulations were observed (all *p*s < .05, uncorrected; Fig. 3C) in both conditions (parafoveal: 2.25, 5.75, and 8 Hz; peripheral: 2.5, 7.25, and 11.25 Hz), which may appear broadly consistent with those observed in the parafoveal condition of Experiment 1. However, phase analysis showed mixed results (Rayleigh tests: *p*s < .05 for only three frequencies, i.e., 2.25 Hz and 8 Hz in the parafoveal condition and 7.25 and 11.15 in the peripheral condition; Figure 3B). As in Experiment 1-2, these patterns were not consistently expressed across individuals (Supplementary Figure S2A), nor stable over time (Supplementary Figure S2B).

**Figure 3.**
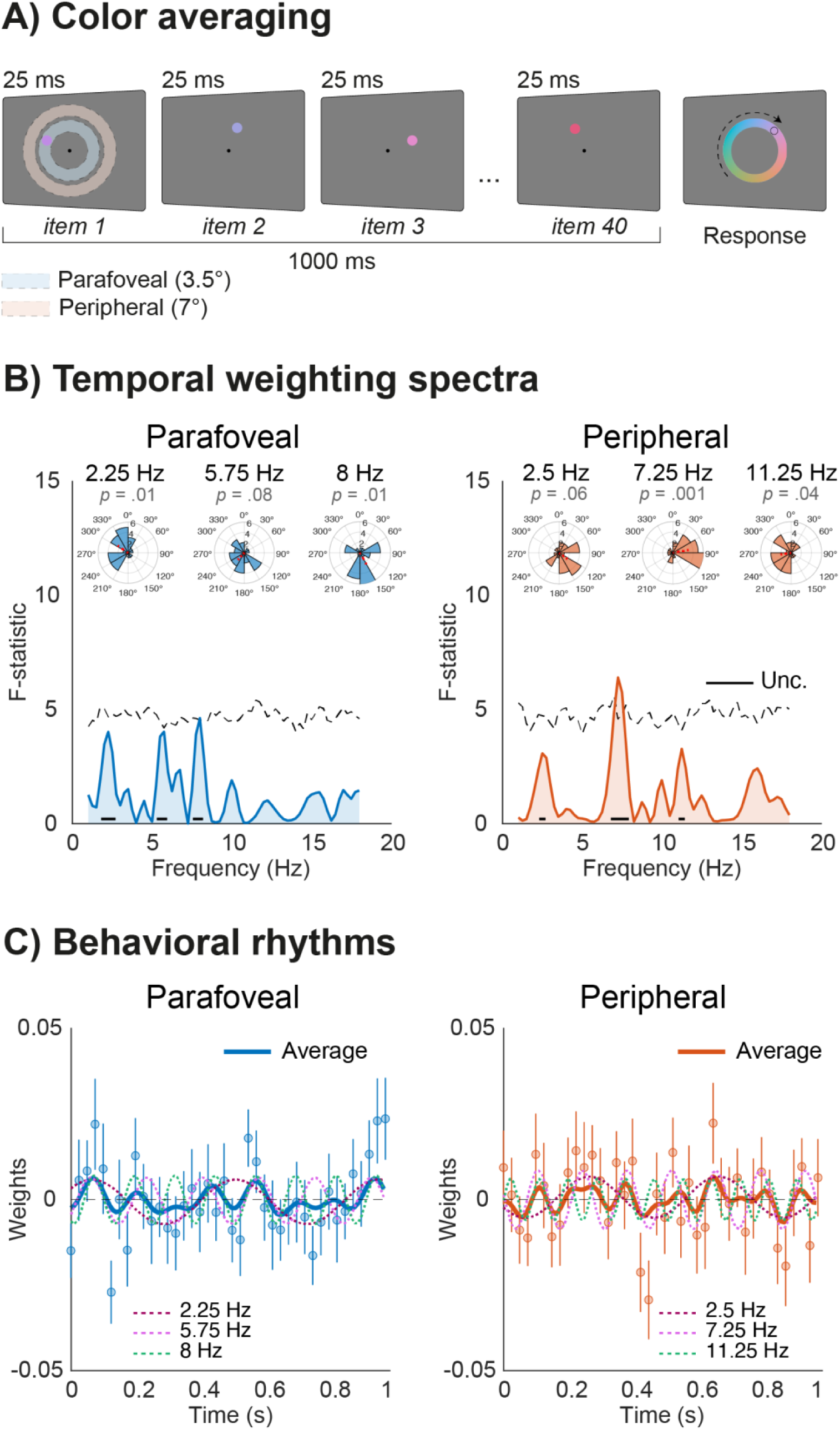
Color judgments. A) In Experiment 3, participants viewed a stream of 40 colored spots presented at 25-ms intervals (40 Hz), either at parafoveal or peripheral eccentricities, and reported the perceived average color by means of a reproduction response. B) Spectral analysis of the temporal weighting profiles in parafoveal (left panel) and peripheral (right panel) conditions. Significant peaks (*p* < .05) are only shown for uncorrected (black lines; GLM F-statistic, see Methods) tests, since no significant peaks were found with FDR-corrected and cluster-based permutation tests. For visualization, the dashed line indicates the 99th percentile of the surrogate distribution. The polar plots show phase clustering for the uncorrected significant peaks, along with the corresponding Rayleigh test p-values. Histograms show the distribution of individual participant phases; the red line indicates the circular mean phase (resultant vector across participants). B) Grand-averaged time course of the temporal weighting profile in parafoveal (left panel) and peripheral (right panel) conditions. Blue and orange lines represent the mean of uncorrected significant rhythms in the parafoveal and peripheral conditions, respectively. Individual uncorrected significant rhythms are also shown for both conditions. Dots and error bars indicate the mean at each time point and the corresponding standard error of the mean (SEM).

## Discussion

In three experiments, we investigated whether perceptual averaging over time exhibits rhythmic structure. We used a weighted-averaging model in which reproduction errors were predicted from deviations of individual stimulus samples from the sequence mean. This yielded temporal weighting profiles reflecting the influence of each sample on the final perceptual estimate. We then analyzed the spectral content of these temporal weighting profiles.

The results revealed a heterogeneous pattern across visual features and viewing conditions. Oscillatory components around ∼3–4 Hz and ∼7–8 Hz emerged during orientation averaging in the parafoveal condition of Experiment 1, with weaker and non-significant trends in Experiment 3 involving color averaging. In contrast, slower rhythmic components around ∼1–1.75 Hz appeared across eccentricity conditions in Experiments 1 and 2, with trends present in Experiment 3 at ∼2–2.5Hz. Overall, these findings indicate that rhythmic fluctuations in temporal weighting depend on visual feature and viewing condition, rather than reflecting a general rhythmic signature.

As mentioned, previous work on behavioral rhythms has relied largely on dense sampling paradigms involving briefly presented, static and isolated stimuli (Landau & Fries, 2012; Fiebelkorn et al., 2013; Song et al., 2014; Drewes et al., 2015; Huang et al., 2015; Michel et al., 2021; Chota et al., 2022), an approach that typically requires large numbers of trials. Here, we adopted a different approach based on perceptual averaging of dynamic stimulus sequences. This offers two main advantages. First, it allows the investigation of behavioral rhythms under conditions that more closely resemble continuous perceptual processing. Second, it substantially increases the amount of information available per trial by estimating temporal weights from the contribution of all stimuli within a sequence, thereby reducing the number of trials required to characterize behavioral rhythms.

That said, our initial application of this approach revealed a heterogeneous pattern, suggesting that rhythmic fluctuations in behavior may be less consistent across perceptual tasks than sometimes implied by general rhythmic-sampling accounts (Landau & Fries, 2012; Samaha & Postle, 2015).

One possibility is that this heterogeneity reflects genuine feature- and task-specific periodic modulations in perceptual averaging, driven by partially distinct underlying mechanisms. For instance, different visual features may rely on partially distinct coding and integration architectures.

Orientation perception is initially supported by orientation-selective channels in early visual cortex (Hubel & Wiesel, 1962; Shapley et al., 2003), whereas size perception may depend more on broader spatial pooling mechanisms that integrate local signals across space (Ariely, 2001; Yildirim et al., 2018). Differences in the temporal dynamics of these computations could therefore produce distinct rhythmic signatures during perceptual averaging. Under this view, the absence of clear rhythmic components in the color experiment may reflect differences in the coding dynamics or temporal integration of color information (Chetverikov et al., 2017; Song et al., 2019; Virtanen et al., 2020) relative to orientation or size. However, we are not aware of direct neurophysiological evidence supporting feature-specific oscillatory mechanisms of this kind, particularly with respect to the differences observed between parafoveal and peripheral conditions in Experiment 1.

An alternative possibility is that the observed heterogeneity reflects the fact that rhythmic components in behavior are generally weak and only become detectable under particular task contexts. In this view, one important factor may be task difficulty, which several studies suggest can modulate both the magnitude and frequency of behavioral rhythms (Dugué et al., 2017; Re et al., 2019; Merholz et al., 2021). We cannot draw firm conclusions regarding the role of difficulty here, as it was not matched across experiments and features. A qualitative observation consistent with this is that orientation and color judgments (Experiments 1 and 3), both involving a circular response space (degrees), differed substantially, with larger response variability in Experiment 3 (Supplementary Figure S1). One might thus speculate that reduced precision in color judgments, possibly reflecting higher task difficulty, may have decreased the sensitivity of our method to detect weak rhythmic components. However, a simple account based on task difficulty cannot explain all within-experiment results. For instance, in Experiment 1, distinct rhythmic components emerged across eccentricity conditions despite comparable behavioral performance. Conversely, in Experiment 2, similar rhythmic components were observed despite differences in performance between conditions. Together, these findings suggest that task difficulty alone is unlikely to fully account for the observed heterogeneity.

A third possibility is that these effects, which here manifested as consistent phase alignment across participants at specific frequencies, are influenced by substantial inter- and intra-individual variability. In line with this, some participants showed stronger alignment with the group mean phase at the group-level peak frequencies than others, and only a minority exhibited high alignment across all significant peaks and conditions, with these effects not being consistently sustained across long blocks of trials (Supplementary Figure S2). This variability complicates the possibility to generalize the characteristics of behavioral rhythms at the group level and suggests that rhythmic structure in performance may instead arise from processing modes that vary in their characteristics and temporal dynamics across and within individuals (Kaltenmaier et al., 2025; Tosato et al., 2025).

The dominant structure of behavioral rhythms observed here falls within a low-frequency regime (1–2.5 Hz) that does not clearly map onto the faster theta- and alpha-range fluctuations emphasized in prior work (Kienitz et al., 2021; Keitel et al., 2022). However, this regime overlaps with accounts of decision formation in which slow delta-band oscillations (∼1–3 Hz) modulate evidence accumulation, leading to phase-dependent fluctuations in the impact of momentary evidence on choice (Wyart et al., 2012). In this view, the low-frequency behavioral structure observed here, characterized by early positive and later negative weighting, may reflect the temporal characteristics of slow evidence accumulation during iterative updating of the estimated average. Such slow push– pull dynamics may lead to asymmetric weighting across sequence positions, in a manner reminiscent of patterns observed in primacy and recency effects (Hubert-Wallander & Boynton, 2015; Do et al., 2022; Yoo et al., 2025), early facilitation and later inhibitory processes (Klein, 2000; Song et al., 2014) and patterns reported in serial dependence, in which perceptual decisions are attracted toward earlier events while being repelled from more recent ones (Pascucci et al., 2019, 2023).

The temporal scale of these effects is also consistent with long-lasting integration windows reported in studies of postdictive and perceptual integration phenomena (Herzog et al., 2020), in which visual information is integrated over hundreds of milliseconds before a conscious percept emerges, with integration windows extending up to approximately 450 ms (∼2.22 Hz) (Menétrey et al., 2023, 2026). It is possible that these slow fluctuations provide a coarse temporal scaffold for integrating visual events and for “making sense” of their spatiotemporal structure (Herzog et al., 2020; Pascucci & Kristjánsson, 2026). Within this framework, faster rhythmic modulations, such as those in the theta and alpha bands typically reported with static and briefly presented stimuli, may be attenuated or averaged out by slower dynamics that dominate evidence accumulation in continuous stimulation contexts. This could help explain why putative rhythmic sampling signatures are generally not consciously experienced as periodic fluctuations under natural viewing conditions, with only rare reports of such phenomenology under specifically tailored experimental contexts (Nakayama et al., 2018).

We do not claim that the slow rhythms observed here, postdictive effects, and decision-related accumulation dynamics reflect the same underlying mechanisms. However, we would like to emphasize this converging evidence across domains, pointing to the observation that slow temporal structure may represent a relevant component of perceptual processing in continuous stimulation, beyond the involvement often attributed to theta- and alpha-band activity.

In sum, we introduce a novel approach for assessing temporal structure in behavioral responses during perceptual averaging. This framework allows the estimation of stimulus-specific contributions within continuous sequences and provides a flexible alternative to traditional cue–target paradigms. At the same time, our findings raise open questions about when rhythmic components emerge, how stable they are across conditions and individuals, and the role of slower fluctuations.

## Methods

### Participants

In total, 75 different participants took part in the experiments (25 in Experiment 1, 12 females; age range 18–30; 25 in Experiment 2, 14 females; age range 19–26; 25 in Experiment 3, 13 females; age range 18–25) and received monetary compensation (25 chf/h). All participants were naive as to the purpose of the experiment and had normal or corrected-to-normal vision, as assessed through the Freiburg acuity test (threshold for inclusion: >1; Bach, 1996). For Experiment 3, red–green vision deficiency was additionally screened using 12 Ishihara pseudoisochromatic plates (threshold for inclusion: <1 error). The study was approved by the local ethics committee (Commission cantonale d’éthique de la recherche sur l’être humain, Canton of Vaud, Switzerland; protocol number: 2024-00863; title: The ebb and flow of human brain activity, cognition and performance) following the Declaration of Helsinki. Written informed consent was obtained from each participant before the experiment.

### Apparatus

The stimuli were presented on a Dell 4K G3223Q monitor (32-inch screen, resolution 3840 × 2160 pixels, refresh rate 120 Hz) and generated using Psychophysics Toolbox version 3.0.19 (Brainard, 1997) running on MATLAB (MathWorks Inc., Natick, MA, USA) version 24.2 (R2024b). Experiments were conducted in a dimly lit room, with participants seated 60 cm away from the screen.

### Stimuli and task procedure

In Experiment 1, participants were presented with streams of Gabor patches defined by a peak contrast of 25% Michelson, a spatial frequency of 2 cycles per degree, and a Gaussian envelope with a standard deviation of 0.375°. The patches appeared sequentially at 12 positions arranged along invisible circular grids. In the parafovea condition, the radius of the circular grid and the center of each Gabor was 3.5° from the center of the screen, whereas in the periphery condition, this distance was 7°. Each Gabor patch was displayed for 25 ms, and consecutive patches never appeared at the same location. Each trial involved a stream of 40 Gabors lasting exactly 1 s. On each trial, the orientation of individual Gabors was sampled from a distribution with a predefined average orientation, randomly determined across the full orientation space, and with a standard deviation of 10°. Participants indicated the perceived average orientation by adjusting a response tool consisting of two dark gray circles connected by an imaginary line, moving the mouse upward or downward to rotate the tool, and confirming their selection with a mouse click.

Experiment 2 used the same locations, timing, and presentation constraints as Experiment 1, but stimuli were light gray filled spots of varying sizes. On each trial, the average size of the spots was determined by randomly sampling from a distribution with a mean of 0.75°, 0.945°, 1.19°, or 1.5° and a standard deviation of 0.2°. Participants reproduced the perceived average size by adjusting a response spot presented at the center of the screen, moving the mouse upward or downward to increase or decrease its size, and confirming the response with a mouse click.

Experiment 3 followed the same presentation parameters as Experiments 1 and 2, but the stimuli were colored spots, with a diameter of 1°. The color of each spot was drawn from a distribution with the average defined by randomly sampling from a circular color wheel in the HCL color space (360 discrete colors) across trials, with a standard deviation of 20. Participants reproduced the perceived average color of the stimulus stream on each trial using a central color wheel response tool and confirmed their selection with a mouse click.

The procedure was similar across all experiments. Each trial began with a fixation period of 500 ms, followed by the stimulus stream. After the 1 s stimulus stream and a blank interval of 500 ms, the response tool appeared. Following the participant’s response, an inter-trial interval with random duration (between 1 and 2 s, sampled in steps of 100 ms) was presented before the next trial. The parafovea and periphery conditions were presented in separate blocks (5 blocks of 80 trials per condition, 10 blocks and 800 trials in total), with self-paced breaks allowed between blocks. Participants completed all blocks of one condition before proceeding to the other condition. The order of conditions was balanced across participants, such that half of the participants began with the parafovea condition and half began with the periphery condition. Participants completed a brief practice session before each experiment and were instructed to maintain fixation at the center of the screen while attending to all elements in the display. Each experiment lasted approximately one hour.

### Temporal weighted average model

Before analysis, adjustment responses were cleaned to remove outliers. In experiments with stimulus features defined in circular space (Experiments 1 and 3), outliers were identified using a von Mises– uniform mixture model (Zhang & Luck, 2008) fit to the individual error distributions, and trials

classified as likely guesses based on the posterior probability of the uniform component (threshold = 0.5) were excluded from further analysis. In the experiment with linear stimulus features (Experiment 2), outliers were defined as responses with errors exceeding 2.5 standard deviations from the individual-specific mean error. Responses faster than 0.2 s or longer than 6 s following the response tool onset were also excluded. Participants were excluded from further analysis if more than 25% of the trials were excluded or if the correlation between true ensemble mean and response was lower than 0.6 (3 participants in Experiment 1, 4 participants in Experiment 2, 0 participant in Experiment 3). In the retained participants, for Experiments 1, 2, and 3, respectively, less than 3.9%, 1.8%, and 2.2% of trials were removed as outliers, and the average correlation between the true ensemble mean and responses exceeded 0.7 (see Supplementary Figure S1).

To estimate the temporal weighted average model, we adapted a method previously used in the context of spatial ensemble perception (Pascucci et al., 2021; Tiurina et al., 2024). Briefly, responses were transformed into errors, and the features of each stimulus (orientation, size, or color, depending on the experiment) were expressed as deviations from the true average feature presented on each trial. These transformed variables approximated a normal distribution centered near 0°. The temporal weighted average model was then estimated using ordinary least squares regression (via the Moore–Penrose pseudoinverse), in which trial-wise errors were modeled as a function of the time series of deviations from the stimulus average. Both errors and deviations were z-scored prior to model estimation. This procedure allowed us to recover the linear weights with which the deviation of each stimulus at each temporal position in the stream contributed to the response error on each trial. The resulting regression coefficients defined a temporal weighting profile reflecting how strongly each stimulus, presented at 25 ms intervals, influenced the perceived average. These time series of weights were estimated separately for each participant and each condition (parafovea and periphery) and subsequently submitted to spectral analysis.

### Spectral analysis

To quantify periodic components in the temporal weighting profiles, we performed a frequency-domain analysis using a general linear model (GLM) approach. For each frequency of interest, the temporal weighting profiles were modeled as a linear combination of sine and cosine functions at that frequency (Tosato et al., 2022; Xie et al., 2025). The temporal weighting profiles were concatenated across participants, and the model was fit using the Moore–Penrose pseudoinverse.

For each frequency, the strength of the rhythmic component was quantified using the F-statistic associated with the model fit, which captures the proportion of variance in the temporal weighting profiles explained by the sinusoidal predictors at that frequency. This procedure yielded, for each condition, a spectrum of F-values.

Statistical inference on the temporal weighting spectra was assessed using both false discovery rate (FDR) correction (Storey, 2002) and cluster-based permutation (Nichols & Holmes, 2002; Maris & Oostenveld, 2007) testing across frequencies. For the FDR approach, frequency-specific p-values derived from the GLM F-statistic were corrected using the Benjamini–Hochberg procedure (α = .05). For the cluster-based analysis, an uncorrected threshold (α = .05) was used to define candidate clusters as contiguous frequencies exceeding the threshold, and cluster mass was computed as the sum of the F-statistic within each cluster. A null distribution was constructed by applying the same spectral analysis to surrogate datasets (N = 1000), in which the temporal order of stimuli within each trial was randomly shuffled, thereby disrupting the correspondence between stimulus sequence and behavioral responses across trials. For each surrogate, the maximum cluster mass across frequencies was retained (Maris & Oostenveld, 2007). Cluster-level p-values were obtained by comparing observed cluster masses to this null distribution.

For visualization, the 99th percentile of the surrogate distribution was computed at each frequency. Peak frequencies were identified based on both FDR-corrected and cluster-corrected results, allowing sensitivity to both narrowband effects (e.g., confined to a single frequency) and broader effects spanning adjacent frequencies. Uncorrected results are also reported for exploratory purposes.

To assess phase consistency across participants at the peak frequencies identified from the F-statistic analysis, we extracted the complex Fourier coefficients at those frequencies from the individual temporal weighting profiles. Phase angles were computed for each participant, and circular statistics were used to quantify phase alignment, or ‘clustering’, across participants. Specifically, for each peak frequency identified in each condition, the circular mean phase was estimated from the complex phase vectors, and phase clustering was quantified using the phase-locking value (Lachaux et al., 1999), defined as the magnitude of the mean resultant vector. Statistical significance of phase clustering was assessed using the Rayleigh test for non-uniformity of circular data (Watson & Williams, 1956). This analysis provided, for each condition and peak frequency, an estimate of the average phase, the degree of phase alignment across participants, and the corresponding statistical significance.

## Acknowledgments

The authors would like to thank Toscane Revillard and Paola Biocchi for their assistance with data collection.

## Data and material availability

Data and analysis scripts supporting the results reported in the main text and supplementary materials will be made publicly available in an online repository upon publication.

## Supplementary Material

**Supplementary Figure 1.**
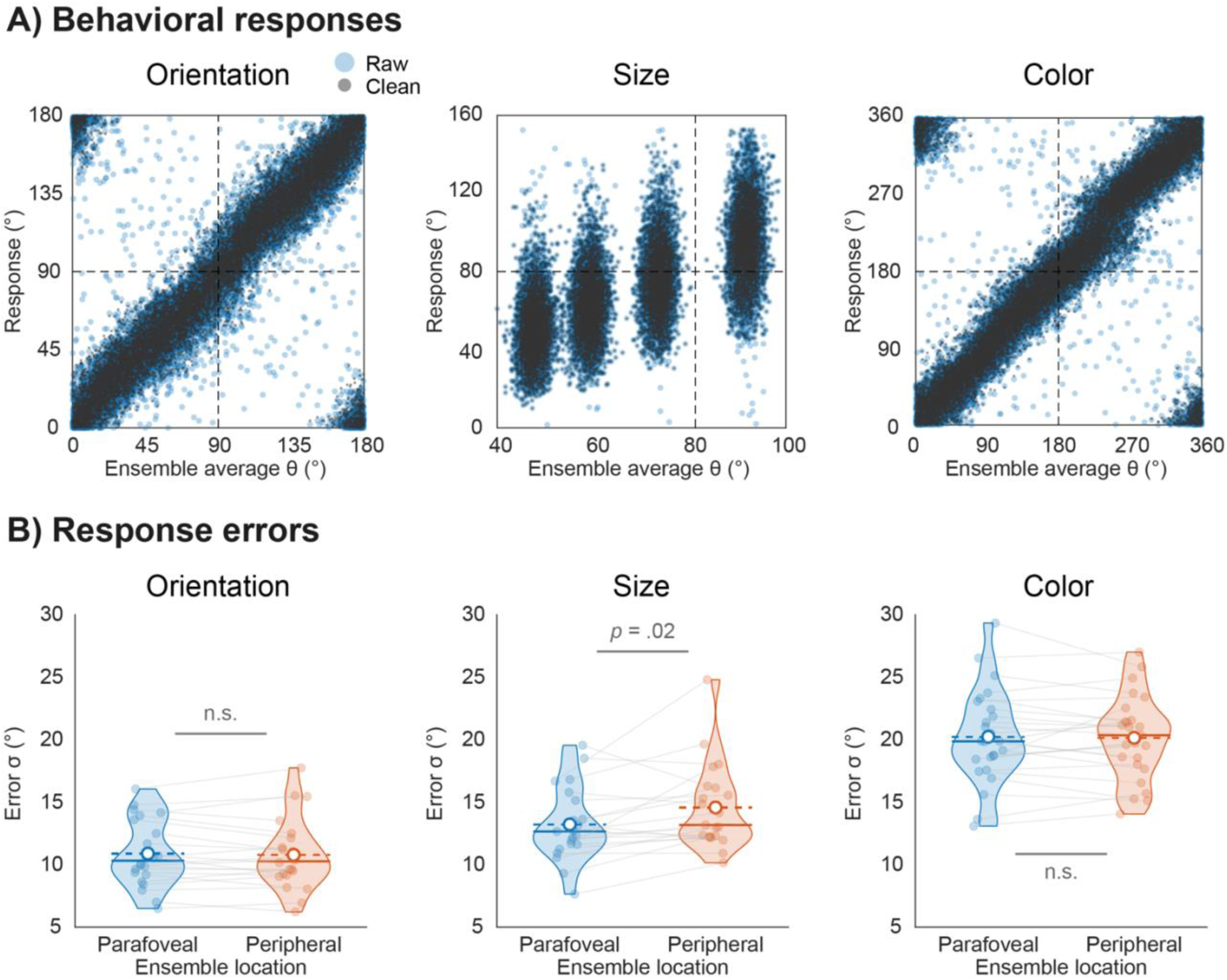
A) Participants’ raw responses (blue dots) and cleaned responses (dark gray; see Methods) plotted as a function of the true ensemble average (Θ, in degrees) for orientation (Experiment 1, left panel), size (Experiment 2, middle panel), and color (Experiment 3, right panel). Data are pooled across parafoveal and peripheral conditions. B) Comparison of the variability of response errors between parafoveal (blue) and peripheral (orange) conditions for orientation (left panel), size (middle panel), and color (right panel) reports. Circular standard deviations are shown for orientation and color, and standard deviations for size. Blue and orange dots indicate individual participants’ mean error, in parafoveal and peripheral conditions, respectively. The white circle and dashed line indicate the group mean after outlier removal, while the solid line indicates the group median. Statistical results are from paired t-test.

**Supplementary Figure 2.**
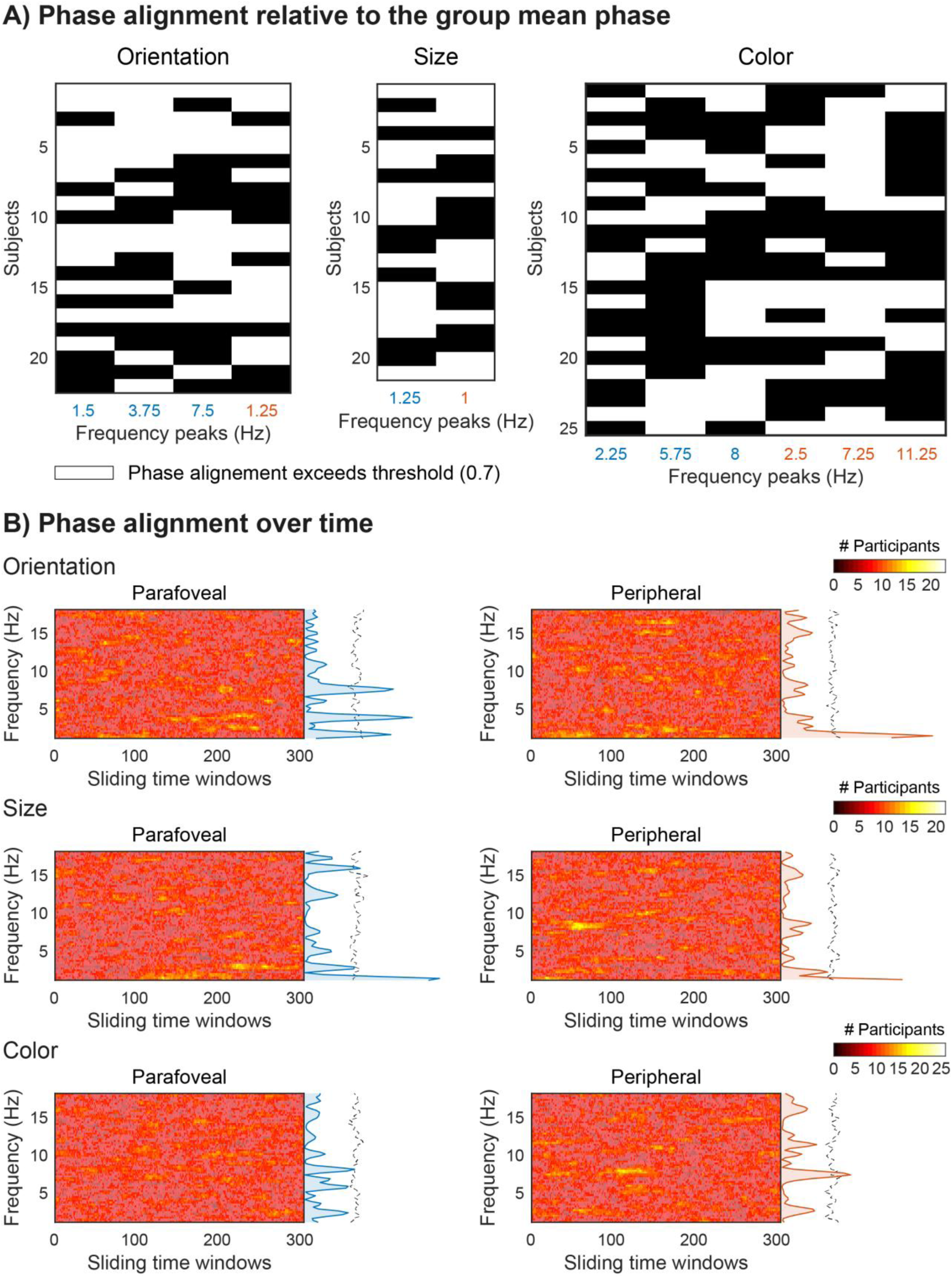
A) Individual phase alignment relative to the group mean phase, shown for each experiment and eccentricity condition at each significant frequency. Each column corresponds to a significant peak frequency identified in the spectral analysis (blue: parafoveal; orange: peripheral), and each row corresponds to a participant. Values indicate whether individual phase estimates at each frequency exceed a predefined alignment threshold (0.70) relative to the circular group mean phase, computed as 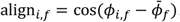, where ϕ_*i,f*_ is the phase of participant *i* at frequency *f*, and 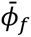 is the circular mean phase across participants. Values above threshold indicate strong phase alignment with the group reference phase. This provides a measure of the consistency with which participants express phase alignment at group-defined rhythmic components. White squares indicate suprathreshold alignment (≥ 0.7), whereas black squares indicate subthreshold alignment. These binarized matrices were then used to estimate the proportion of participants showing strong alignment for each peak frequency, as well as across all significant frequencies identified in the main analyses (see main text). B) Time-resolved group-level phase alignment derived from sliding-window regression (100-trial windows). For each participant and condition, behavioral data were modeled using a sliding GLM, as in the main analysis (see Methods), yielding time-resolved regression weights that were transformed into the frequency domain using a fast Fourier transform. Phases of the resulting complex spectral coefficients were extracted per participant and frequency bin, and aligned to the circular group mean phase. Values reflect the number of participants exceeding a predefined phase-alignment threshold (0.70) at each frequency and sliding time window, providing a time-resolved measure of cross-participant consistency in phase structure. For reference, spectral results of the temporal weighting profiles for parafoveal and peripheral conditions (computed across all trials; see Figures 1B, 2B, and 3B) are shown on the right of each panel, together with the 99th percentile of the surrogate distribution.

## References

Anderson, N. H. (1967). Application of a weighted average model to a psychophysical averaging task. Psychonomic Science, 8(6), 227–228. 10.3758/BF03331634

Ariely, D. (2001). Seeing Sets: Representation by Statistical Properties. Psychological Science, 12(2), 157–162. 10.1111/1467-9280.00327

Bach, M. (1996). The Freiburg Visual Acuity Test—Automatic Measurement of Visual Acuity: Optometry and Vision Science, 73(1), 49–53. 10.1097/00006324-199601000-00008

Bonnefond, M., Jensen, O., & Clausner, T. (2024). Visual Processing by Hierarchical and Dynamic Multiplexing. Eneuro, 11(11), ENEURO.0282-24.2024. 10.1523/ENEURO.0282-24.2024

Brainard, D. H. (1997). The Psychophysics Toolbox. Spatial Vision, 10(4), 433–436.

Brookshire, G. (2022). Putative rhythms in attentional switching can be explained by aperiodic temporal structure. Nature Human Behaviour, 6(9), 1280–1291. 10.1038/s41562-022-01364-0

Ceylan, G., Herzog, M. H., & Pascucci, D. (2021). Serial dependence does not originate from low-level visual processing. Cognition, 212, 104709. 10.1016/j.cognition.2021.104709

Chetverikov, A., Campana, G., & Kristjánsson, Á. (2017). Representing Color Ensembles. Psychological Science, 28(10), 1510–1517. 10.1177/0956797617713787

Chota, S., Leto, C., Van Zantwijk, L., & Van Der Stigchel, S. (2022). Attention rhythmically samples multi-feature objects in working memory. Scientific Reports, 12(1), 14703. 10.1038/s41598-022-18819-z

Do, J., Eo, K. Y., James, O., Lee, J., & Kim, Y.-J. (2022). The Representational Dynamics of Sequential Perceptual Averaging. The Journal of Neuroscience, 42(6), 1141–1153. 10.1523/JNEUROSCI.0628-21.2021

Drewes, J., Zhu, W., Wutz, A., & Melcher, D. (2015). Dense sampling reveals behavioral oscillations in rapid visual categorization. Scientific Reports, 5(1), 16290. 10.1038/srep16290

Dugué, L., Xue, A. M., & Carrasco, M. (2017). Distinct perceptual rhythms for feature and conjunction searches. Journal of Vision, 17(3), 22. 10.1167/17.3.22

Fiebelkorn, I. C., & Kastner, S. (2019). A Rhythmic Theory of Attention. Trends in Cognitive Sciences, 23(2), 87–101. 10.1016/j.tics.2018.11.009

Fiebelkorn, I. C., Saalmann, Y. B., & Kastner, S. (2013). Rhythmic Sampling within and between Objects despite Sustained Attention at a Cued Location. Current Biology, 23(24), 2553–2558. 10.1016/j.cub.2013.10.063

Fries, P. (2015). Rhythms for Cognition: Communication through Coherence. Neuron, 88(1), 220–235. 10.1016/j.neuron.2015.09.034

Helfrich, R. F., Fiebelkorn, I. C., Szczepanski, S. M., Lin, J. J., Parvizi, J., Knight, R. T., & Kastner, S. (2018). Neural Mechanisms of Sustained Attention Are Rhythmic. Neuron, 99(4), 854–865.e5. 10.1016/j.neuron.2018.07.032

Herzog, M. H., Drissi-Daoudi, L., & Doerig, A. (2020). All in good time: Long-lasting postdictive effects reveal discrete perception. Trends in Cognitive Sciences, 24(10), 826–837.

Huang, Y., Chen, L., & Luo, H. (2015). Behavioral Oscillation in Priming: Competing Perceptual Predictions Conveyed in Alternating Theta-Band Rhythms. The Journal of Neuroscience, 35(6), 2830–2837. 10.1523/JNEUROSCI.4294-14.2015

Hubel, D. H., & Wiesel, T. N. (1962). Receptive fields, binocular interaction and functional architecture in the cat’s visual cortex. The Journal of Physiology, 160(1), 106–154. 10.1113/jphysiol.1962.sp006837

Hubert-Wallander, B., & Boynton, G. M. (2015). Not all summary statistics are made equal: Evidence from extracting summaries across time. Journal of Vision, 15(4), 5. 10.1167/15.4.5

Kaltenmaier, A., Davis, M. H., & Press, C. (2025). Fixed and flexible perceptual rhythms. Trends in Cognitive Sciences, 29(10), 903–913. 10.1016/j.tics.2025.05.011

Keitel, C., Ruzzoli, M., Dugué, L., Busch, N. A., & Benwell, C. S. Y. (2022). Rhythms in cognition: The evidence revisited. European Journal of Neuroscience, 55(11–12), 2991–3009. 10.1111/ejn.15740

Kienitz, R., Schmid, M. C., & Dugué, L. (2021). Rhythmic sampling revisited: Experimental paradigms and neural mechanisms. European Journal of Neuroscience, ejn.15489. 10.1111/ejn.15489

Klein, R. M. (2000). Inhibition of return. Trends in Cognitive Sciences, 4(4), 138–147. 10.1016/S1364-6613(00)01452-2

Lachaux, J.-P., Rodriguez, E., Martinerie, J., & Varela, F. J. (1999). Measuring phase synchrony in brain signals. Human Brain Mapping, 8(4), 194–208. https://doi.org/10.1002/(SICI)1097-0193(1999)8:4%3C194::AID-HBM4%3E3.0.CO;2-C

Landau, A. N., & Fries, P. (2012). Attention Samples Stimuli Rhythmically. Current Biology, 22(11), 1000–1004. 10.1016/j.cub.2012.03.054

Maris, E., & Oostenveld, R. (2007). Nonparametric statistical testing of EEG- and MEG-data. Journal of Neuroscience Methods, 164(1), 177–190. 10.1016/j.jneumeth.2007.03.024

Menétrey, M. Q., Herzog, M. H., & Pascucci, D. (2023). Pre-stimulus alpha activity modulates long-lasting unconscious feature integration. NeuroImage, 278, 120298. 10.1016/j.neuroimage.2023.120298

Menétrey, M. Q., Herzog, M. H., & Pascucci, D. (2026). Sequential neural dynamics underlie unconscious integration and conscious perception of visual stimuli. PLOS Biology, 24(7), e3003894. 10.1371/journal.pbio.3003894

Merholz, G., Grabot, L., VanRullen, R., & Dugué, L. (2021). *Periodic attention operates faster during more complex visual search* [Preprint]. Neuroscience. 10.1101/2021.09.22.460906

Michel, R., Dugué, L., & Busch, N. A. (2021). Distinct contributions of alpha and theta rhythms to perceptual and attentional sampling. European Journal of Neuroscience, ejn.15154. 10.1111/ejn.15154

Nakayama, R., Motoyoshi, I., & Sato, T. (2018). Discretized Theta-Rhythm Perception Revealed by Moving Stimuli. Scientific Reports, 8(1), 5682. 10.1038/s41598-018-24131-6

Nichols, T. E., & Holmes, A. P. (2002). Nonparametric permutation tests for functional neuroimaging: A primer with examples. Human Brain Mapping, 15(1), 1–25. 10.1002/hbm.1058

Ozkirli, A., & Pascucci, D. (2023). State-dependent serial dependence in perceptual decisions (p. 2023.10.19.563128). bioRxiv. 10.1101/2023.10.19.563128

Pascucci, D., & Kristjánsson, Á. (2026). Spatiotemporal routines in visual perception. Nature Reviews Psychology, 1–13. 10.1038/s44159-026-00568-9

Pascucci, D., Mancuso, G., Santandrea, E., Della Libera, C., Plomp, G., & Chelazzi, L. (2019). Laws of concatenated perception: Vision goes for novelty, decisions for perseverance. PLOS Biology, 17(3), e3000144. 10.1371/journal.pbio.3000144

Pascucci, D., Ruethemann, N., & Plomp, G. (2021). The anisotropic field of ensemble coding. Scientific Reports, 11(1), 8212. 10.1038/s41598-021-87620-1

Pascucci, D., Tanrikulu, Ö. D., Ozkirli, A., Houborg, C., Ceylan, G., Zerr, P., Rafiei, M., & Kristjánsson, Á. (2023). Serial dependence in visual perception: A review. Journal of Vision, 23(1), 9. 10.1167/jov.23.1.9

Re, D., Inbar, M., Richter, C. G., & Landau, A. N. (2019). Feature-Based Attention Samples Stimuli Rhythmically. Current Biology, 29(4), 693–699.e4. 10.1016/j.cub.2019.01.010

Samaha, J., & Postle, B. R. (2015). The Speed of Alpha-Band Oscillations Predicts the Temporal Resolution of Visual Perception. Current Biology, 25(22), 2985–2990. 10.1016/j.cub.2015.10.007

Shapley, R., Hawken, M., & Ringach, D. L. (2003). Dynamics of Orientation Selectivity in the Primary Visual Cortex and the Importance of Cortical Inhibition. Neuron, 38(5), 689–699. 10.1016/S0896-6273(03)00332-5

Song, A., Faugeras, O., & Veltz, R. (2019). A neural field model for color perception unifying assimilation and contrast. PLOS Computational Biology, 15(6), e1007050. 10.1371/journal.pcbi.1007050

Song, K., Meng, M., Chen, L., Zhou, K., & Luo, H. (2014). Behavioral Oscillations in Attention: Rhythmic α Pulses Mediated through θ Band. The Journal of Neuroscience, 34(14), 4837– 4844. 10.1523/JNEUROSCI.4856-13.2014

Storey, J. D. (2002). A direct approach to false discovery rates. Journal of the Royal Statistical Society: Series B (Statistical Methodology), 64(3), 479–498. 10.1111/1467-9868.00346

Tiurina, N. A., Markov, Y. A., Whitney, D., & Pascucci, D. (2024). The functional role of spatial anisotropies in ensemble perception. BMC Biology, 22(1), 28. 10.1186/s12915-024-01822-3

Tosato, T., Dumas, G., Rohenkohl, G., & Fries, P. (2025). Performance modulations phase-locked to action depend on internal state. iScience, 28(1). 10.1016/j.isci.2024.111691

Tosato, T., Rohenkohl, G., Dowdall, J. R., & Fries, P. (2022). Quantifying rhythmicity in perceptual reports. NeuroImage, 262, 119561. 10.1016/j.neuroimage.2022.119561

VanRullen, R. (2016). Perceptual Cycles. Trends in Cognitive Sciences, 20(10), 723–735. 10.1016/j.tics.2016.07.006

Virtanen, L. S., Olkkonen, M., & Saarela, T. P. (2020). Color ensembles: Sampling and averaging spatial hue distributions. Journal of Vision, 20(5), 1. 10.1167/jov.20.5.1

Watson, G. S., & Williams, E. J. (1956). On the Construction of Significance Tests on the Circle and the Sphere. Biometrika, 43(3/4), 344. 10.2307/2332913

Wyart, V., de Gardelle, V., Scholl, J., & Summerfield, C. (2012). Rhythmic Fluctuations in Evidence Accumulation during Decision Making in the Human Brain. Neuron, 76(4), 847–858. 10.1016/j.neuron.2012.09.015

Xie, X.-Y., Burr, D. C., & Morrone, M. C. (2025). Recent, but not long-term, priors induce behavioral oscillations in peri-saccadic vision. Communications Psychology, 3(1), 41. 10.1038/s44271-025-00224-7

Yildirim, I., Öğreden, O., & Boduroglu, A. (2018). Impact of spatial grouping on mean size estimation. *Attention, Perception*, & Psychophysics, 80(7), 1847–1862. 10.3758/s13414-018-1560-5

Yoo, M., Bahg, G., Turner, B., & Krajbich, I. (2025). People display consistent recency and primacy effects in behavior and neural activity across perceptual and value-based judgments. *Cognitive, Affective*, & Behavioral Neuroscience, 25(4), 923–940. 10.3758/s13415-025-01285-1

Zhang, W., & Luck, S. J. (2008). Discrete fixed-resolution representations in visual working memory. Nature, 453(7192), 233–235. 10.1038/nature06860

